# The framework of lncRNAs and genes at early pollen developmental stage in a PTGMS wheat line

**DOI:** 10.1101/2020.06.05.136606

**Authors:** Jian-fang Bai, Zi-han Liu, Yu-kun Wang, Hao-yu Guo, Li-Ping Guo, Zhao-guo Tan, Shao-hua Yuan, Yan-mei Li, Ting-ting Li, Wen-jing Duan, Jie-ru Yue, Feng-ting Zhang, Chang-ping Zhao, Li-ping Zhang

## Abstract

Wheat photo-thermosensitive genic male sterile (PTGMS) line is a vital material in the two-line hybrid wheat breeding system in which functional pollen production is highly associated with temperature during early developmental stage. Understanding the potential mechanism of pollen infertility induced by low temperature in PTGMS wheat is crucial for the effective utilization of genetic resources to guide wheat breeding. Herein, we combined full-length single-molecular sequencing and Illumina short reads sequencing data to obtain the high-resolution spatio-temporal transcriptome map of pollen under low temperature stress at mother cell, dyad and tetrad stages in PTGMS line BS366. Cytological descriptions and whole transcriptome analysis revealed a global landscape of low temperature altered pollen fertility transformation via regulating the transcriptional patterns of cytoskeleton-related lncRNAs and their target genes, which involved in the calcium signaling and vesicle trafficking pathways on cytoskeleton homeostasis at different stages of meiosis. Overall, our results provided the transcriptional and cytological evidences for understanding the low temperature-induced pollen sterility deficiency in PTGMS wheat line.

## Introduction

Pollen abortion contains multiple physiological, biochemical, and molecular changes such as the abnormal degradation of the tapetum (Zheng et al., 2019), defective pollen wall (Marianne et al., 2002; Wu et al., 2015), the level of kinetic of ATPases (Sane et al., 1997), the distribution and concentration of Ca^2+^ in anther (Tian et al., 1998), the regulation of the cytoskeleton (Tang et al., 2012; Wang et al., 2018), accumulation ROS in tapetum (Liu et al., 2018a), and the abnormities of cell signaling transductions (Liu et al., 2018b). Based on mentions above, it is well known that cytological biological events and related genes may play important roles in regulation of plants male fertility. In CMS lines wheat, premature or delayed PCD by tapetal cells disorganized the supply of the nutrients to microspores, thereby resulting in pollen abortion (Meng et al., 2016). Previous studies suggested this irregular tapetal PCD was tightly controlled by evolutionarily conserved transcriptional cascades (Liu et al., 2020). In recent years, hybrid breeding has a remarkable success in several allogamous species such as maize, sunflower, sorghum, sugar beet, and rye, but not be fully exploited in autogamous crops (Longin et al., 2012). Wheat is an autogamous crop, however, hybrid seed production requires cross-pollination of the female parent by pollen from the male parent. Photoperiod and/or thermo-sensitive genic male sterility (P/TGMS) is an important material in two-line breeding system to explore the potential of heterosis. Previous studies showed that the male sterility of P/TGMS lines are contributed to abnormal pollen development (Bai et al., 2017). Further cytological studies showed that P/TGMS wheat, it exhibited disordered distribution of the cytoskeleton, including microflaments and microtubules when exposed to a sterile environment during the fertility-sensitive stage (Tang et al., 2011; Wang et al., 2018). Additionally, previous RNA-seq studies also have mostly focused on the transcripts of male sterility and thousands of differentially expressed genes have been reported (Liu et al., 2016). As we all know, in male sterile wheat, the process of pollen abortion reflects extremely complex reprogramming of gene expression involving chromatin modification, transcription, posttranscriptional processing, posttranslational modification, and protein turnover. However, the complex posttranscriptional and translational levels molecular mechanisms of the male sterility, especially the wheat P/TGMS line induced by low temperature, are currently still not clear.

As a hexaploid, wheat has a large and complex genome, estimated to reach approximately 17G, which composes three closely-related and independently maintained genomes that are the result of a series of naturally occurring hybridisation events. With the continuous advancement of technology, second-generation sequencing technology (e.g., Illumina sequencing), could feature high-throughput capability and provide high-quality reads. However, the short-read length potentially introduces errors in inaccurately identify the transcript results. Third-generation sequencing is a single-molecule real-time sequencing technology, (e.g., PacBio sequencing), could provide full-length transcript information, detect single-molecule structure, provide complete mRNA structure and is, therefore, well suited for transcript recovery and isoform detection in species with well sequenced and/or incomplete genome sequences (Abdel-Ghany et al., 2016; Wang et al., 2016). In some crop plants, many scientists have been studied the isoform change when plant response to stress at gene transcriptional levels, such AS level and long non cording RNA (lncRNA) (Abdel-Ghany et al., 2016; Wang et al., 2016) using combination of TGS and NGS technology. AS is a widely recognized RNA processing mechanism in eukaryotic species, playing a major role in the molecular biology of the cell, and within humans it has been implicated in multiple genetic disorders (Wang et al., 2016). AS is a critical posttranscriptional event which comes from alternate splice site choices in a single gene locus, including intron retention (IR), exon skipping (ES), alternative 5’ splicing site (Alt5’SS) and alternative 3’ splicing site (Alt3’SS) (Wang et al., 2019). In higher plant, it has been reported that the AS events are involved in a wide range of developmental and physiological processes including responses to stress. For example, about 60% of Arabidopsis intron-containing genes are generated by AS (Marquez et al., 2012). It has been demonstrated that AS is important for cold response when SFs mis-expressed during cold sensitivity or tolerance treatment (Laloum et al., 2018). In wheat, Liu et al. (2018) performed genome-wide analysis of alternative splicing (AS) responses to drought stress (DS), heat stress (HS) and their combination (HD) in wheat seedling to investigate the regulation of AS during these stress processes (Liu et al., 2018c). More recently, Wang et al. (2019) also investigated the spatio-temporal landscape of heat adaptations in wheat filling grain and flag leaves at transcriptional and AS levels by hybrid sequencing (second-and third-generation sequencing). These studies strongly suggest that AS networks are central co-ordinators of the stress response. Although AS plays an important role in stress response, it is not clear in studies related to pollen abortion. In addition to AS the lncRNA were also generated by hybrid sequencing (Wang et al., 2019). Many studies have shown that lncRNA could regulate genes at the transcriptional and post-transcriptional levels by acting as signals, decoys, scaffolds, and guides (Heo and Sung, 2011). However, virtually nothing is known about the extent and timing of the contribution of AS and lncRNA or how AS and lncRNA determine the dynamic changes of transcriptome required for regulating male sterility for P/TGMS line. Here, to illustrate the regulation of these factors and genes on fertility transformation, we used hybrid sequencing strategy and used a conventional fertile wheat variety as a control and differential genes background to analyze the role of AS and lncRNA in regulating male sterility during fertility transformation stages in wheat PTGMS line BS366.

## Results

### Morphological characteristics of mature pollen under different conditions

In order to identify the influence of environment on the fertility of PTGMS line BS366, we observed the phenotype of BS366 at the trinucleate stage under different fertility conditions and used conventional wheat Jing411 as control (Fig. 1). Scanning electron microscopy (SEM) examination at trinuclear stage revealed that epidermis cells of the Jing411 in two conditions and BS366 anthers in fertile conditions were arranged closely, on the contrary, the epidermis cells of BS366 anthers in sterile conditions were incomplete and occurred loss. Moreover, the inner epidermal ubisch bodies of anther for BS366 in sterile conditions were abnormal and accumulated more sparsely distributed. Based on observations of microspores at the trinucleate stage, the microspores were uniformly spheroid and had finely reticulate ornamentation on their surface in anther of Jing411 and BS366 in fertile conditions, while the sterile microspores were extremely atrophied. According to I2-KI staining, BS366 in fertile conditions pollen grains were 60% stained black, whereas pollen grains were all wrinkled and inadequately stained in sterile condition, and exhibited completely aborted characteristics, indicating that low temperature could induce the male fertility conversion in PTGMS line BS366.

**Figure 1:**
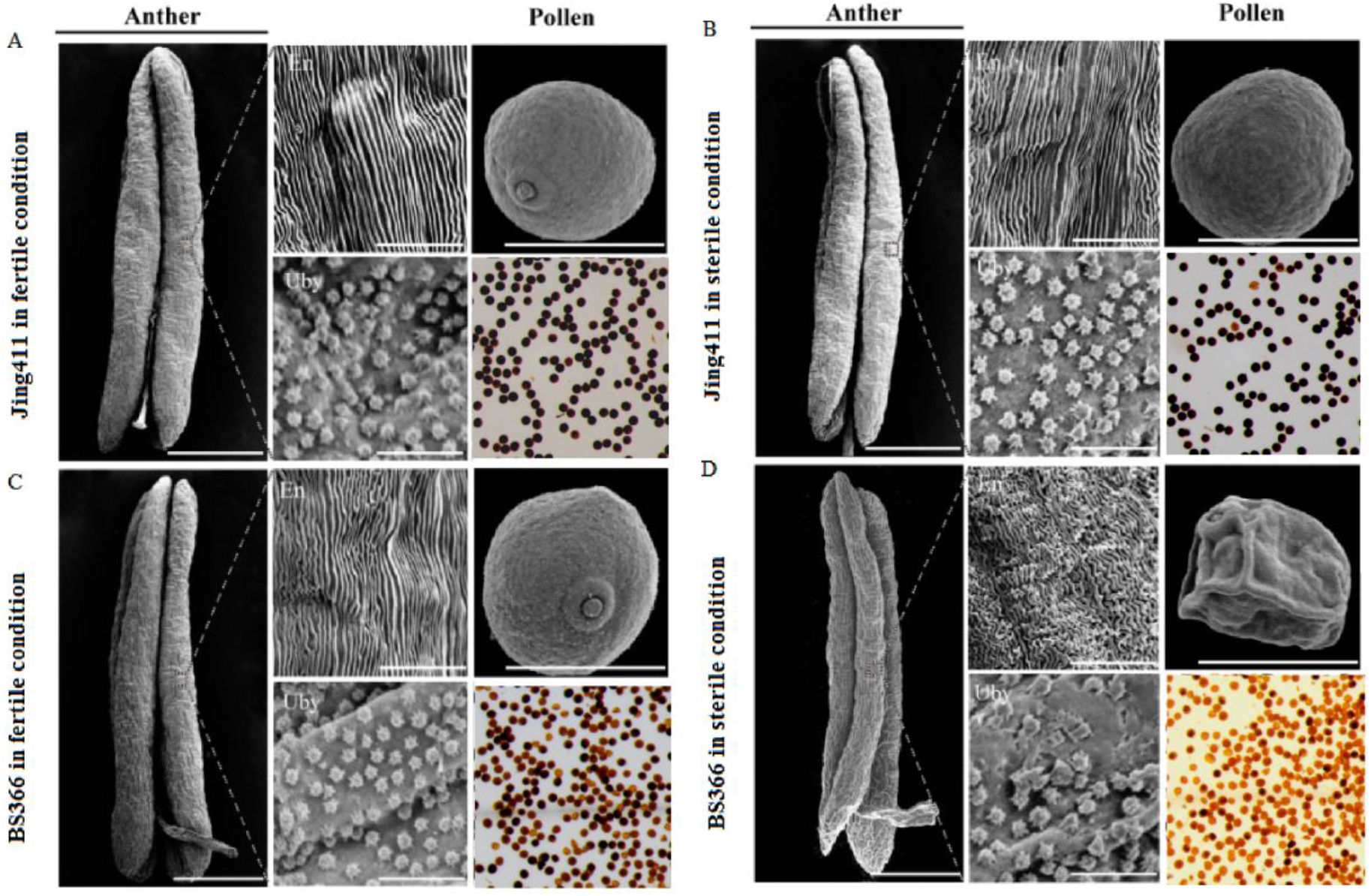
Phenotypes of mature anthers and pollen of Jing411 (**A, B**) and BS366 (**C, D**) at the trinucleate stage under fertile and sterile conditions. Scale bars in anther are equivalent to 1 mm, in epidermis, ubisch bodies and pollen are equivalent to 50μm. Abbreviations: epidermis (E), ubisch bodies (Uby).

### Overview of sequencing data

Previous study showed that the period from the pollen mother cell stage to the tetrad stage is the most sensitive to low temperature for the pollen of BS366 (Bai et al., 2017). And previous cytological studies also showed that there were abnormalities of film-forming body and cell plate in this process (Tang et al., 2011). To comprehensively investigate the wheat transcriptomes present during the fertility transition, hybrid sequencing were performed on the anther of BS366 and Jing411 from different fertility condition with three pollen development stages (S1: pollen mother cell stage, S2: dyad stage, and S3: tetrad stage) (Fig. S1). Totally, 247,486 and 240,993 circular consensus sequence reads for BS366 and Jing411 were yielded, respectively (Table S2). In total, 209,967 and 190,280 reads of full-length non-chimeric (FLNC) (84.84% and 78.96%) from these circular consensus sequence reads, were identified based on the inclusion of 5’and 3’ primer, and 3’ poly(A) tails, followed by error correction. Then, 209,967 and 190,280 high-quality isoforms were uniquely mapped to the IWGSC RefSeq v1.0 (Table S3). Finally, 16,000 and 14,864 transcripts were obtained from BS366 and Jing411, followed by error correction using Illumina short reads (Table S4). The length of transcripts was mainly concentrated in the range of 103-104 for BS366, Jing411 and IWGSC RefSeq v1.0, suggesting the generated data are accurate and reliable, could be used as a reference in this study (Fig. 2 and Fig. S2). These transcripts were derived from 13,577 and 13,010 gene loci. Of which, 1,185 and 902 are new gene loci and 6,248 and 4,918 are new transcripts (Table S4). By comparison with the IWGSC RefSeq v1.0 annotation, PacBio transcripts could be classified into seven groups including PacBio data, DEG, DAS, lncRNA, fusion gene analysis (Fig. 3).

**Figure 2:**
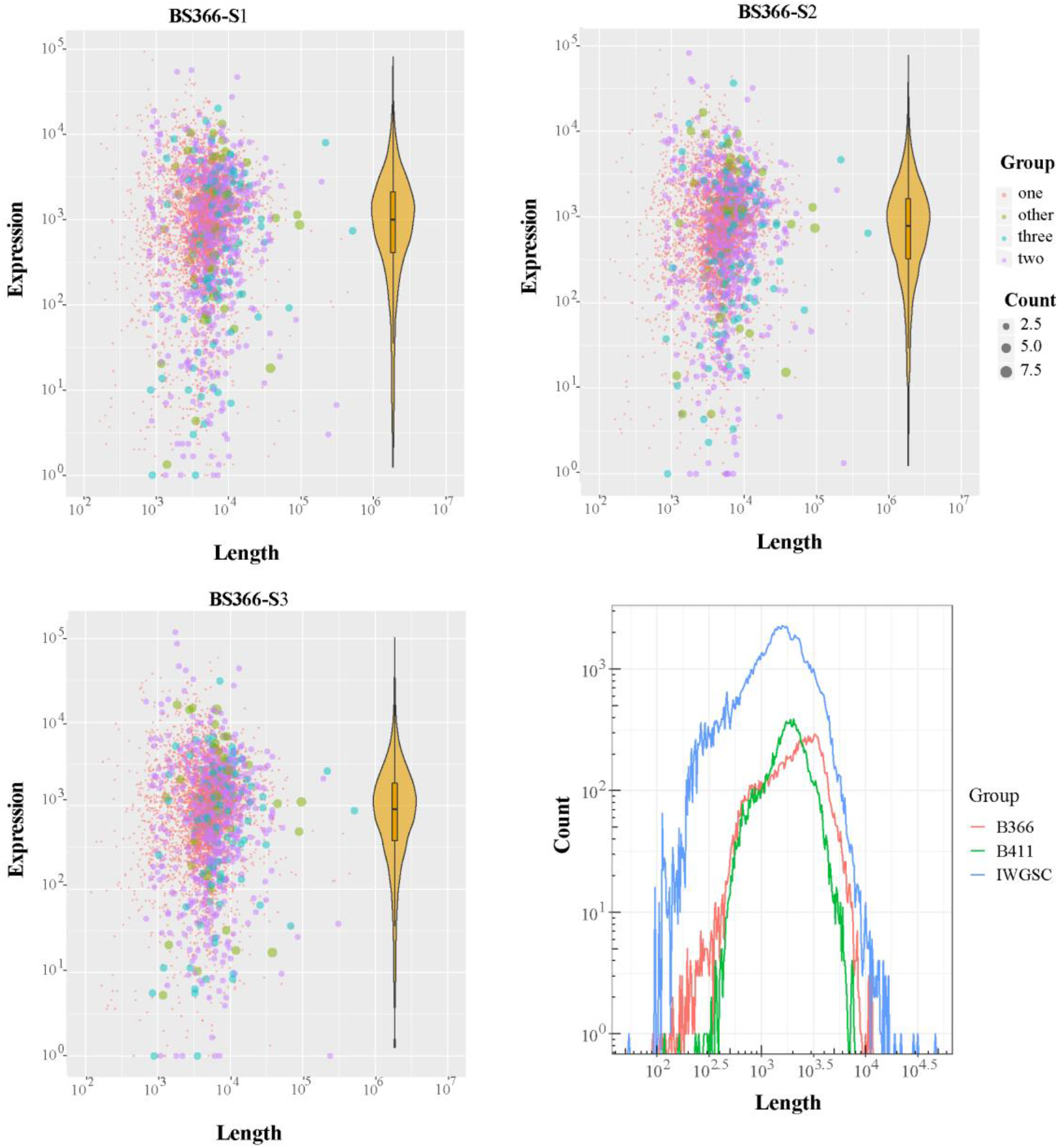
Summary of the direct RNA sequencing data of BS366. **A-C:** The bubble scatter plots show the relationship between the fraction of detected transcripts by the direct RNA sequencing with the transcript length and the level transcript expression. The violin-boxplots on the right show the overall distribution of the expression of transcripts. **D:** The histogram plot shows the distribution of read length of high quality reads obtained from BS366 (red), Jing411 (green) and IWGSC, respectively.

**Figure 3:**
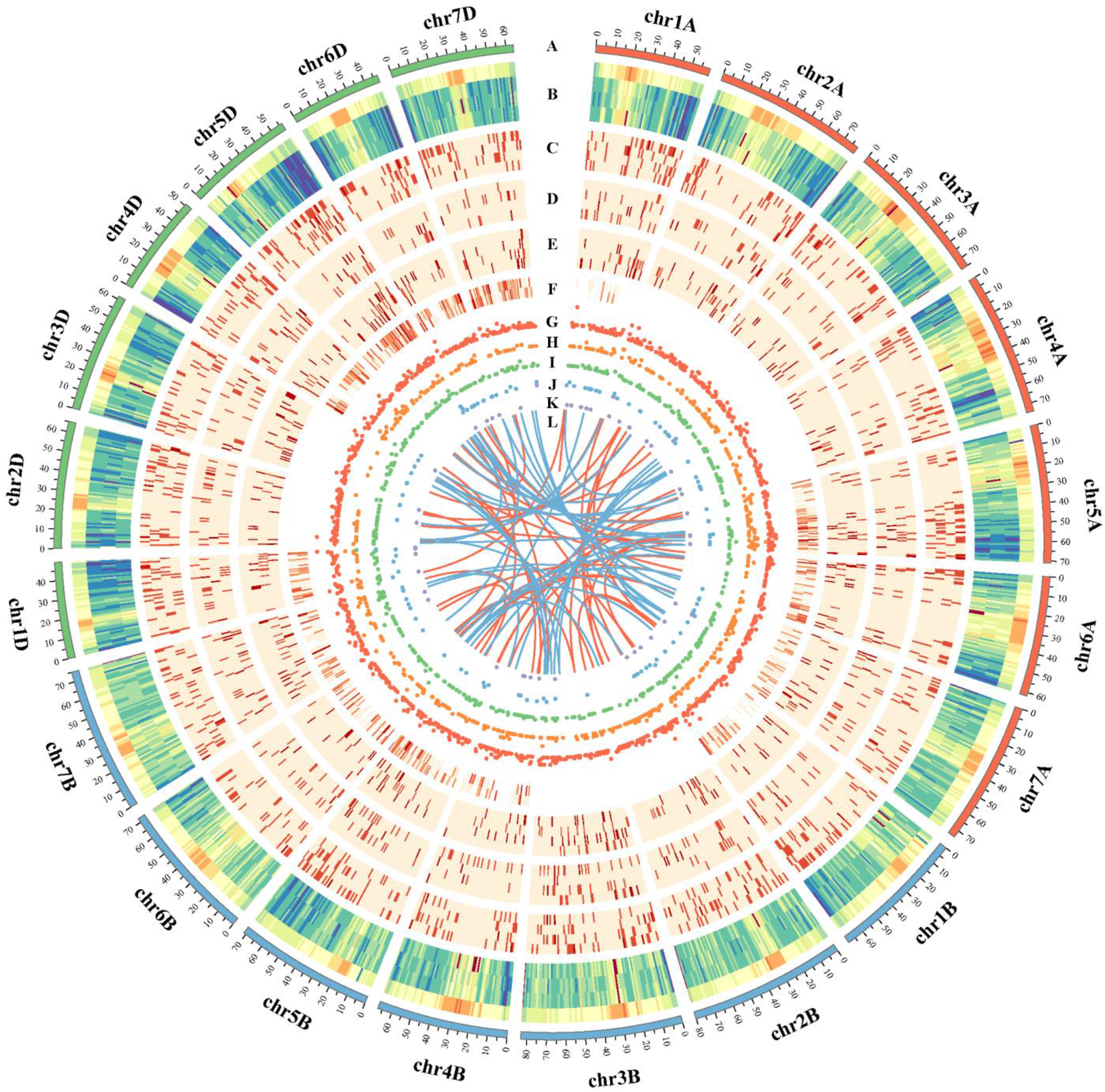
CIRCOS visualization of different data at the genome-wide level. The density was calculated in a 10-Mb sliding window **A:** Karyotype of the wheat genome. **B:** Comparison of transcript density between the IWGSC RefSeq v1.0 annotation and the PacBio data. From the upper to lower tracks: transcripts in IWGSC RefSeq v1.0, transcripts in BS366 and 411, transcripts in BS366 and 411 in pollen, respectively. **C-E:** Distribution of differentially expressed genes (DEGs) and differentially spliced genes (DSG) for S3, S2 and S1 of BS366 and Jing411 in between fertile and sterile conditions. From the upper to lower tracks in each part: DEGs for BS366 in between fertile and sterile conditions, DSGs for BS366 in between fertile and sterile conditions, DEGs for Jing411 in between fertile and sterile conditions, DSGs for Jing411 in between fertile and sterile conditions. **F:** Distribution of transcription factors in BS366 (uper track) and Jing411 (lower track). **G-K:** Identificated of lncRNAs from pfam (**G**), CPC (**H**), CPAT (**I**), CNCI (**J**) and overlap of them (**K**). **L:**Linkage of fusion transcripts in BS366(red) and Jing411(blue).

### Identification of alternative splicing events during fertility transition

AS is an important biological event of post-transcriptional regulation in organisms (Wang et al., 2019). A few studies have explored the relationship between AS and fertility transition which regulated by external environment in plants (Capovilla et al., 2015). Therefore, we analyzed the patterns of fertility transition-induced AS events from the RNA sequencing data. In this study, four main AS events including Alternative 3’ splice site (Alt3’SS), Exon skipping (ES), Intron retention (IR) and Mutually exclusive exon (MEX) were identified from three pollen developments stages of BS366 and Jing411. As shown in Table 1, total 35,248 and 33,099 AS events with 20,312 and 19,490 AS genes were identified from three pollen developments in BS366 and Jing411, respectively. Of which 11,619, 11,840 and 11,789 AS events in BS366 and 10,980, 11,084 and 11,035 AS events were determined on subgenomes A, B and D, respectively (Table S5). It was found that ES was the most abundant (91.08%−91.98%) AS events in both BS366 and Jing411, followed by MEX (4.84%−5.24%), IR (3.03%−3.67%) in BS366 (Table 1), and some orders in Jing411 and for AS genes (Table S5).

### Identification of sterility -related AS events and genes in BS366

The high-resolution temporal transcriptomes allowed us to determine the specific stage at which differentially spliced genes (DSGs) and differentially expressed genes (DEGs, fold change ≥2.0, FDR-adjusted *P*-value <0.05) showed a significant change, along with the magnitude and trend of that change. To identify the specific time of significant changes of DSGs and DEGs as well as analysis the sterility-related DSGs and DEGs in BS366, DSGs and DEGs from three key pollen development stages of BS366 during fertility transition BS366 were screened. By comparing the DSGs of three stages, in total, 108, 130 and 141 genes with 118, 141 and 160 AS events were identified from pollen mother cell stage, dyad stage and tetrad stage respectively after critically filtering process. For DEGs, in total, 39, 73 and 412 DEGs were identified from three stages (Figure 4A). The change trend of DSGs and DEGs in the three stages was synergistically rising, in which the DSGs and DEGs contained in tetrad stage have significant changes, which may be the key stage of fertility transformation.

**Figure 4:**
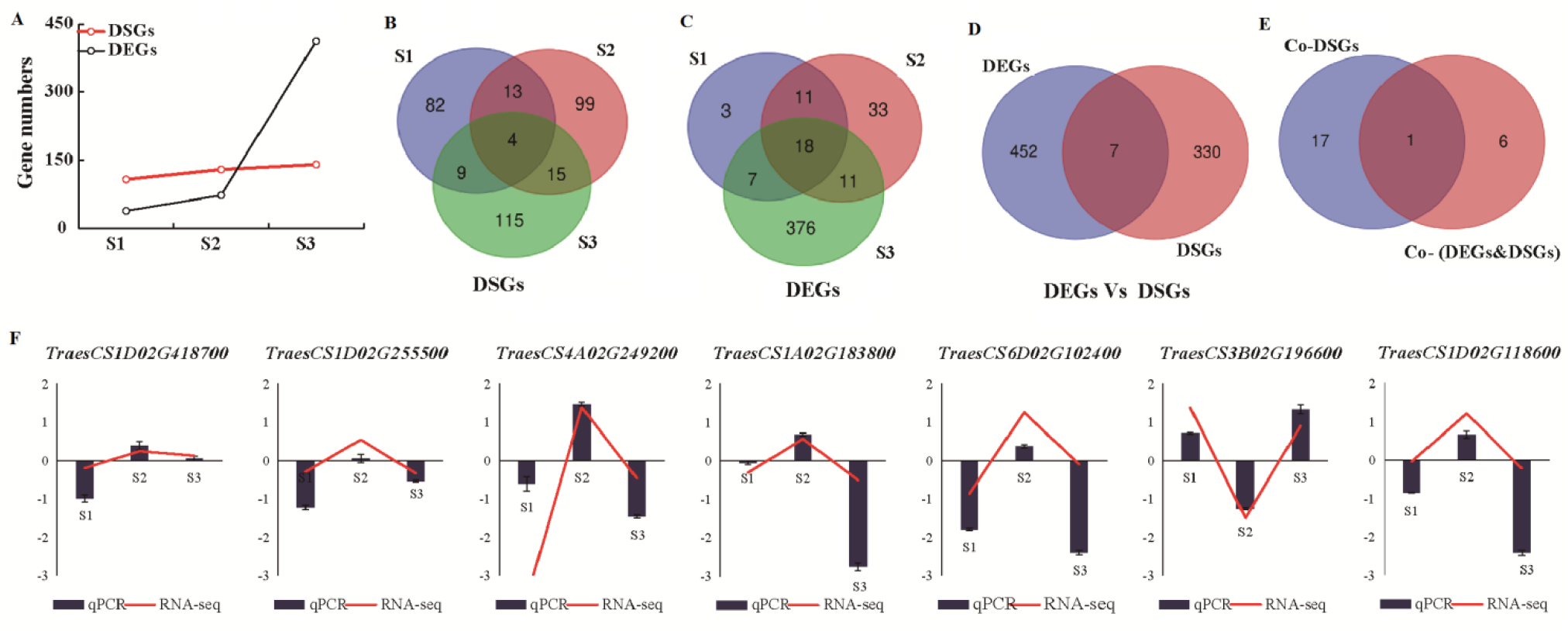
Identification and comparison analysis of sterility-related AS genes and sterility-related genes during fertility transition. **A:** The changes of gene number of DSGs and DEGs during anther development stages. **B-E:** Venn diagram of DSGs (**B**) and DEGs (**C**) in three stages, genes in DSGs and DEGs (**D**), and genes in common DSGs and common DSGs &DEGs (**E**) **F:**qPCR analysis of seven genes of common DSGs &DEGs

Here four DSGs were found in shared DSGs and DEGs in the three stages (Fig. 4B), in which the genes encoding zinc finger domain protein 1A (*TraesCS5D02G371100*) and UTP--glucose-1-phosphate uridylyltransferase (*TraesCS5A02G353700*) involved in pollen formation (Chivasa et al., 2013), and the gene encoding coatomer subunit gamma-2 (*TraesCS1D02G156000*) was associated with vesicle transport (Hamlin et al., 2014). Moreover, cytoskeleton-related gene kinesin-4 (*TraesCS3B02G196600*), vesicle transport-related gene encoding vacuolar protein sorting-associated protein (*TraesCS4B02G382900*), and pollen formation-related genes encoding zinc finger BED domain-containing protein (*TraesCS3B02G126800*) and MYB transcription factors (*TraesCS3B02G243600*) were found in co-DEGs of the three stages (Figure 4C). Interestingly, the gene encoding kinesin-4 (*TraesCS3B02G196600*) was differentially expressed in three stages at the same time and occurred differentially AS in stage 3 (Fig. 4D and E), and the kinesin-4 has been proved to be involved in vesicle transport and cytoskeleton formation (van Riel et al., 2017), thereby inferring that AS of this gene could work together with transcriptional regulation to involve in fertility transformation of PTGMS line BS366. qPCR analysis verified the correctness of RNA-seq results (Fig. 5F). The primers used in this study were listed in Table S1.

**Figure 5:**
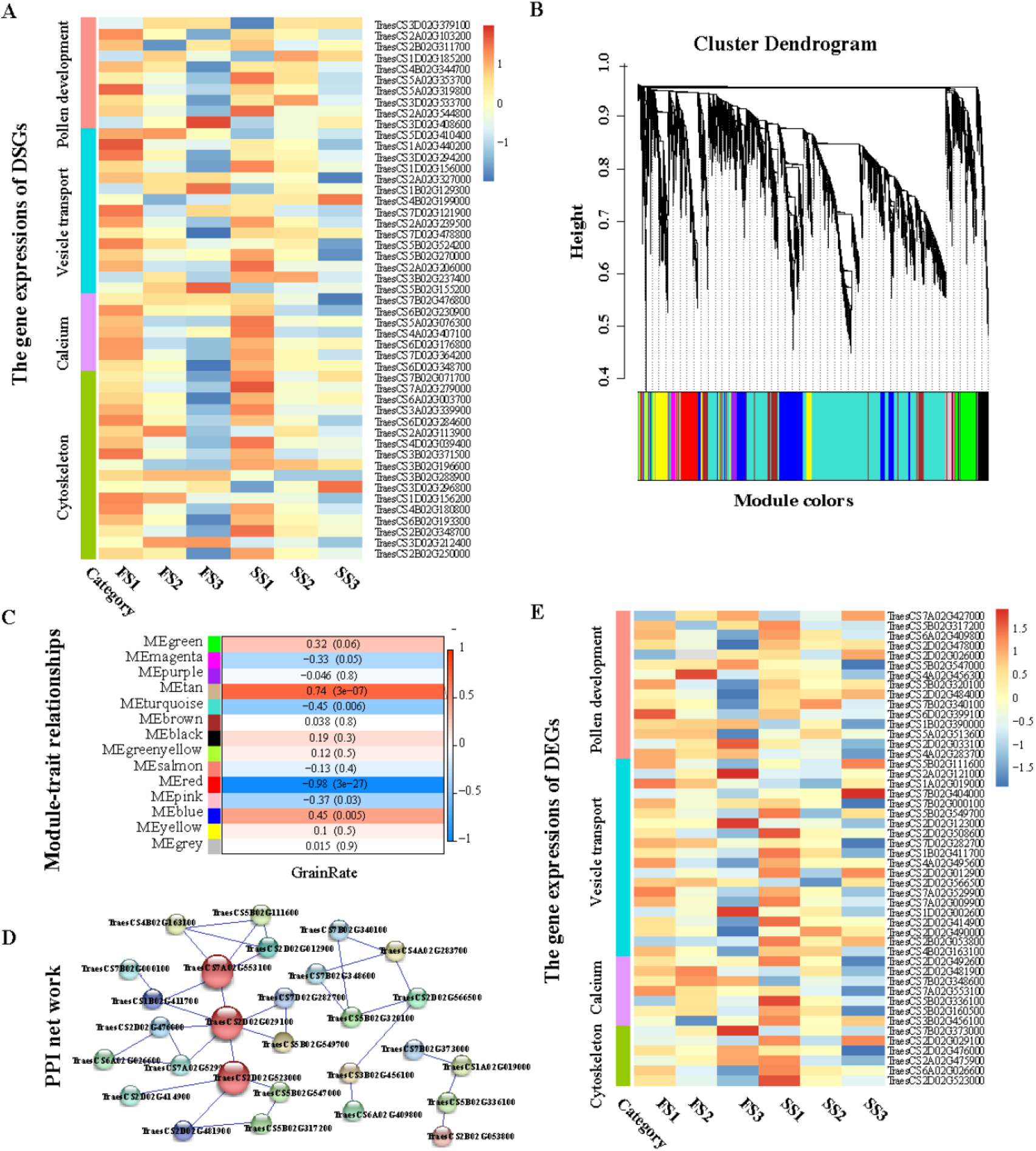
Analysis male sterility-related DSGs and DEGs. **A:**Heat map for pollen sterility-related DSGs. **B:** Hierarchical cluster tree showing the modules of co-expressed genes, where the lower panel shows the Modules in different colors. **C:** Module-trait correlations and corresponding *p*-values (inparentheses), where the left panel shows the module eigen genes and the right panel shows a color scale for the module trait correlations ranging from −1 to 1. **D:** Cytoscaper epresentation of the co-expressed genes in important pathways in the red module. **E:** Heat map for male sterility-related DEGs.

### Comparative analysis of the biological functions regulated at AS and transcription levels

GO analysis showed that the most of DSGs were correlated with some cytological and molecular events in the process of pollen development including cytoskeleton (such as “cytoskeleton” and “cytoplasmic microtubule organization”), calcium regulation (such as “calcium ion binding” and “calcium-dependent phospholipid binding”), vesicle transport (such as “vesicle-mediated transport” and “vacuole”), and pollen formation (such as “cell wall organization” and “glucose-1-phosphate uridylyltransferase activity”) during the three key stages of fertility conversion (Table S6). It has been reported these GO terms identified are involved in the male sterility, for example, the DSGs encoding the protein MOR1 (*TraesCS3B02G371500* in S1, *TraesCS3A02G339900* in S2, and *TraesCS3B02G371500* in S3) was critical for the orderly assembly of microtubules (Kawamura et al., 2006). Microtubules and microfilaments control the whole double fertilization process of pollen, is necessary for the normal development of pollen. Therefore, low temperature induced DSGs was a crucial further layer of regulation for pollen development, thereby possibly leading to fertility conversion.

To further identify the sterility-related genes, weighted gene co-expression network analysis (WGCNA) were performed with all DEGs from BS366 based on the seed setting rate (Table S7). The analysis of module-trait relationships analysis showed that the module ‘Red’ (*r* = −0.98, *p* = 3e-27) was highly correlated with male sterility in the six samples (Fig. 5B and C). Interestingly, GO analysis suggested that these DEGs in this module were also mainly concentrated in the GO terms associated with cytoskeleton, calcium regulation, vesicle transport, and pollen formation, which were consistent with that of DSGs. Furthermore, cytoskeletal-regulatory complex EF hand (*TraesCS7A02G553100*), actin-related protein 9 (*TraesCS2D02G029100*), and microtubule-associated protein RP/EB family member 3 (*TraesCS2D02G523000*) were greater connectivity in these DEGs, suggesting they may be strongly associated with male sterility (Fig. 5D). Thus, transcriptional regulation may play a major role in cytoskeleton, calcium regulation, vesicle transport, and pollen formation, which coordinates with AS regulation to induce pollen abortion in wheat.

### Differences of diverse transcription factors (TFs) during fertility transition

Transcription factors (TF) that perceive environmental signals and activate the expression of related genes play master roles in gene regulatory networks in the processes of growth and development including pollen development in plants (Wang et al., 2018). In this work, total 522 TFs with 633 transcripts from Pacbio data were annotated which belonging to major 48 families (Table S8). Furthermore, 34 wheat TFs were differentially expressed (DE-TFs), and 6 genes were differentially spliced (DS-TFs) during fertility transition (S1, S2 and S3) in BS366 after removing background, respectively (Fig. 6 and Tables S9). In addition, enrichment analysis was performed on these TFs at each pollen development stage to revel the fertility transition-related signaling. As shown in Fig. 6, bZIP family was significantly enriched in DE-TFs at dyad stage, MYB and bHLH families were enriched in DE-TFs at tetrad stage. More, there were several members of the HMG, ARF, NAC and bHLH families were differentially expressed at the tetrad stage providing evidence that these families participated in fertility transition signaling transduction.

**Figure 6:**
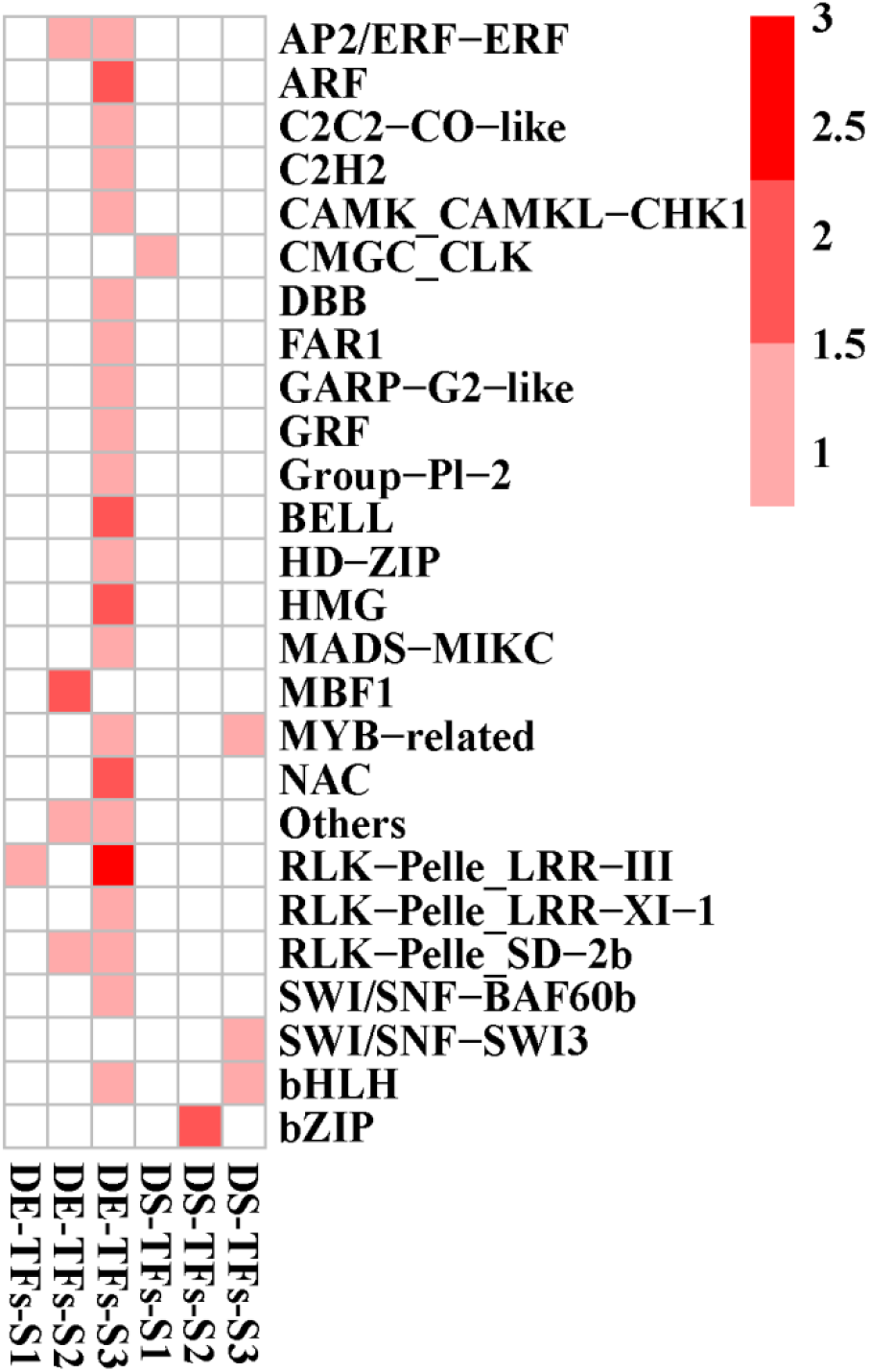
Differentially expressed TFs (DE-TFs) and differentially spliced TFs (DS-TFs) in different anther development stages

### Function analysis of predicted lncRNAs and their targets

After filtering by CPC, CNCI, CPAT and Pfam, total 53 lncRNAs (>200bp) were obtained from Pacbio data (Fig. 7A). Generally, lncRNAs regulate gene expression via *cis* (regulation of neighboring loci) or *trans*-acting mechanisms. It has been proposed that lncRNAs that are synthesized at a low level are likely to act in *cis*, whereas those accumulate at a higher level are able to act in *trans* (Kornienko et al., 2013). Identification and analysis of candidate target genes could provide insight into the functions of lncRNAs in fertility transition of PTGMS line. In this study, 38 ncRNA–mRNA pairs were identified as *cis*-regulation and five were *trans*-regulation lncRNA–mRNA pairs from 53 lncRNAs after filtering (Table S10). In this five *trans*-regulation lncRNA– mRNA pairs, only PB.18919.1 have *trans*-regulation targets and other four lncRNAs have both *cis* and *trans-*regulation targets. There were 14 lncRNAs which no targets were found (Table S10). In addition, like other no cording RNA, such as miRNA (Bai et al., 2017), the same lncRNA can regulate multiple mRNA genes, different lncRNA molecules can also be synergistic regulation of the same mRNA gene (Table S10), indicating that lncRNAs with their targets may participate in multiple regulation pathway.

**Figure 7:**
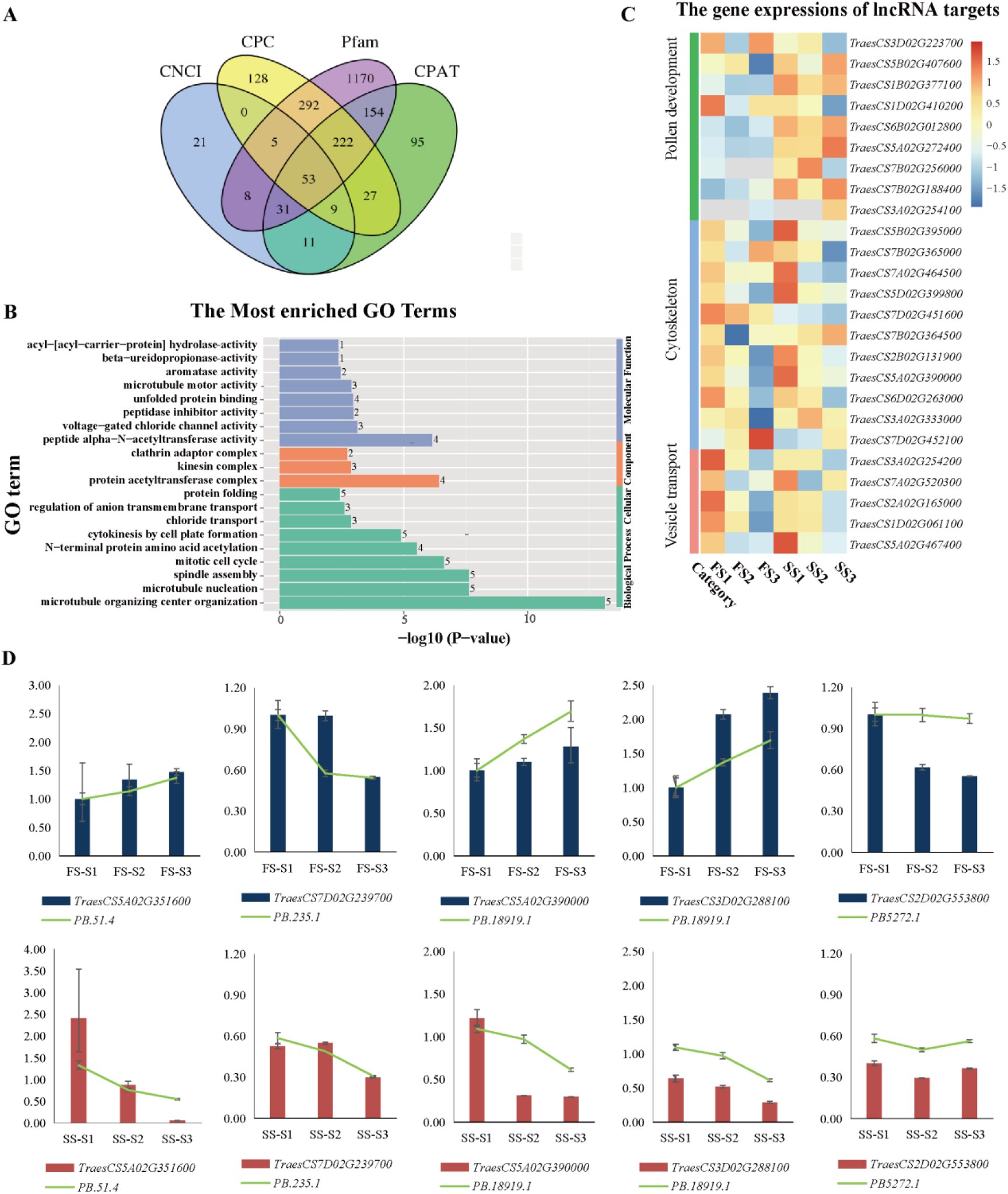
Analysis of identified lncRNAs. **A:** Identified lncRNAs from Pacbio data by using CPC, CNCI, CPAT and Pfam. **B:** Go enrichment analysis of targets of lncRNAs. **C:** Heat map for male sterility-related targets of lncRNA. **D:** qPCR analysis the expression of randomly selected lncRNAs and targets.

To investigate lncRNA functions in regulation of fertility transformation, GO analysis on predicted targets was performed. As shown in Fig. 7B, the most frequent “molecular function” term was “peptide alpha−N−acetyltransferase activity”, followed by “voltage-gated chloride channel activity”, and “microtubule motor activity” for targets. The most frequent “biological process” term was “microtubule organizing center organization”, followed by “microtubule nucleation” and “spindle assembly”. The results indicated that these lncRNA with their targets are involved in process of cell division and important for male sterility in wheat.

qPCR and heatmap analysis were also performed in lncRNAs and their corresponding targets to verified their expression patterns during different pollen development stages (Fig. 7C and D). Coordinated expression was found between most lnRNAs and their targets (Figure 7D and Table S11). PB.51.4, PB.235.1, PB.18919.1 and PB5272.1 with targets (*TraesCS5A02G351600, TraesCS7D02G239700, TraesCS5A02G390000* and *TraesCS2D02G553800* down regulated during pollen development in both fertile condition and sterile condition. However, the other target gene of PB.18919.1, *TraesCS3D02G288100* (Copper transport protein ATX1) showed up-regulated in sterile condition and down-regulated in fertile condition (Fig. 7D).

### RT-PCR validation of the DSGs

In this study, four DSGs were selected randomly from pollen mother cell stage, dyad and tetrad stage in different condition to validate the accuracy of AS events using reverse transcription polymerase chain reaction (RT)-PCR (Fig. 8). The isoforms of each DSG were designed primers to amplify all predicted transcripts and cloned using Sanger sequencing (Table S1). The results as shown by a gel banding pattern in Figure 8, the size of each amplified fragment was consistent with that of predicted fragment (Fig. 8). It was also found that expression of transcript isoforms exhibits a stage-preferential pattern. For example, *TraesCS3B02G326100* (encoding a Lipid-A-disaccharide synthetase protein), which was produced by an A5’SS event, was preferentially expressed at dyad stage and tetrad stage in fertile condition and at pollen mother cell stage in sterile condition. It therefore is example of stage specific RNA isoforms.

**Figure 8:**
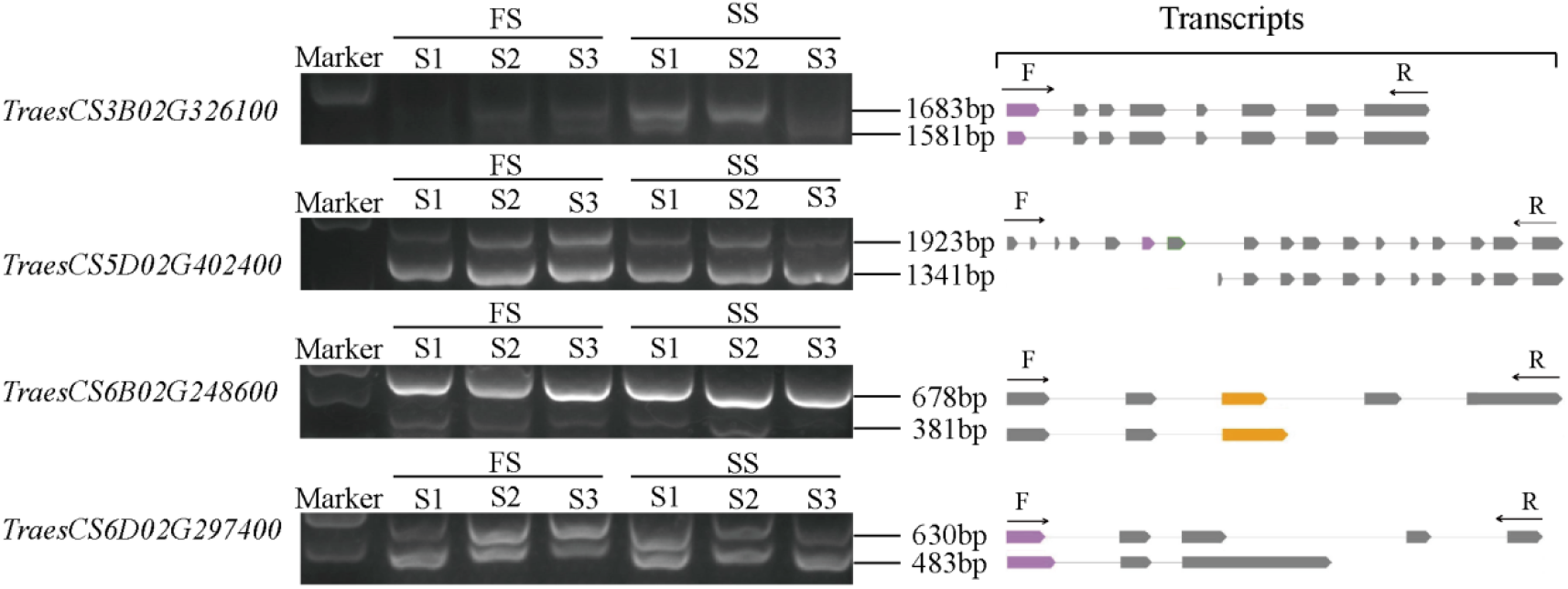
Validation of full-length isoforms using Semi-quantitative RT-PCR. RT-PCR validation of AS events for three genes. Gel bands in each figure show DNA makers and PCR results in three stages under two condition. Transcript structure of each isoform is shown in right panel. Yellow boxes show exons and lines with arrows show introns. PCR primers (F, forward and R, reverse) are shown on the first isoform of each gene. The length of each full-length isoform is shown after the transcript structure.

### Effects of altered alternative splicing and gene expression on important cellular events

Hybrid sequencing showed that most of DSGs, DESs and lncRNA regulated targets could be related to chromosomal movement, process of cell division, cytoskeleton activity, cell plate formation. To further verify the results of the above analysis, the changes of microtubules, microfilaments, cell plate, chromosomes and calcium throughout critical periods of fertility transformation in pollen cells of BS366 under fertile and sterile conditions were observed (Fig. 9A). During pollen mother cell stage, in the fertile condition, the pollen mother cell took on a normal oval shape, the cytoskeleton was evenly distributed and the polar perinuclear microtubules were initially formed. However, in the sterile condition, the overall cellular morphology of pollen mother cell was wrinkled, chromatin was abnormally condensed and arranged scattered in the nuclear region, and microtubules and microfilaments appear as a radial and disordered filament and the polar microtubules were not obvious. Up to dyad stage, the fertile pollen mother cell proceeded normal cytokinesis and formed a distinct cell plate, while the division of sterile cell underwent disruption, and it’s worth noting that the sterile dyad occurred absence of the cell plate. Especially during the tetrad stage, the sterile tetrad happened severe malformation and the cytoskeleton was sparse and disordered compared with BS366 of fertile condition. Thus, under low temperature stress, cytoskeleton related genes and lncRNA regulated targets underwent differential alternative splicing and differential expression, which led to the abnormal meiosis of pollen mother cells, including the concentration of chromatin, the scattered distribution of cytoskeleton and the absence of cell plates.

**Figure 9:**
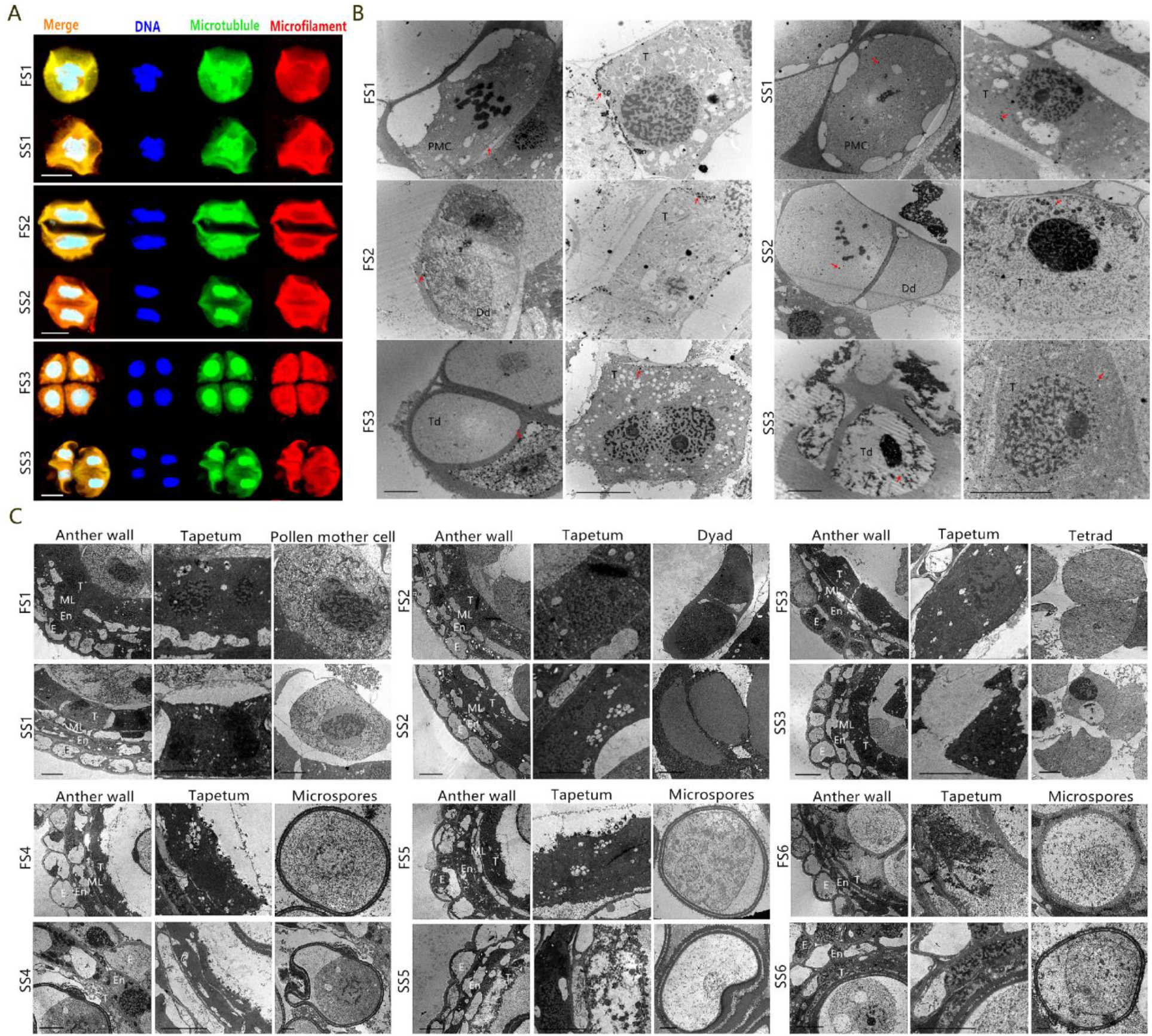
Cytological observation of BS366 under different conditions. **A:** the distribution of cytoskeleton of different conditions from pollen mother stage to terad stage. **B:** the distribution of Ca^2+^ of BS366 under different conditions from pollen mother stage to terad stage. **C:** the ultrastructural observation of anther, tapletum and pollen cell of BS366 under different conditions from pollen mother stage to trinucleate stage. FS1: pollen mother cell stage of fertile condition, FS2: dyad stage of fertile condition, FS3: tetrad stage of fertile condition, FS4: uninucleate stage of fertile condition, FS5: binucleate stage of fertile condition, FS6: trinucleate stage of fertile condition, SS1: pollen mother cell stage of sterile condition, SS2: dyad stage of sterile condition, SS3: tetrad stage of sterile condition, SS4: uninucleate stage of sterile condition, SS5: binucleate stage of sterile condition, SS6: trinucleate stage of sterile condition. Dd: dyad, E: epidermis, En: endothecium, PMC: pollen mother cell, T: tapetum, Td: tetrads. Bars are 4 μm in A and 1μm in B and C.

It was also found that some DEGs such as *TraesCS5B02G336100* (phospholipase D delta), *TraesCS3B02G456100* (60S ribosomal protein L38) and *TraesCS5B02G160500* (calmodulin-binding receptor-like cytoplasmic kinase 3) were all up-regulated and associated with Ca^2+^ distribution (Fig. 5E). Meanwhile, six DSGs encoding calmodulin related protein also was screened (Fig. 5A). Many studies have shown that the dynamic distribution of Ca^2 +^ was related to pollen sterility in anthers. In order to further verify whether fertility conversion is related to Ca^2+^ distribution, we used potassium antimontate to observe the distribution of Ca^2+^ in pollen and tapetum of BS366 under fertile and sterile conditions during pollen development (Fig. 9B). During the process of pollen mother cell division to tetrad, abundant Ca^2+^ precipitates were gradually accumulated on the cell surface, and the distribution of Ca^2+^ in the cytoplasm was very little, which maintained a low concentration of Ca^2+^ distribution environment. However, in the sterile condition, Ca^2+^ was less on the surface of sterile pollen cells, and excessive accumulation of Ca^2+^ precipitated in cytoplasm. Therefore, we concluded that the up-regulation of calcium-related DEGs and the regulation of DSGs may lead to the abnormal function of calcium pump or calcium channel in the cells and the inability to discharge the excess Ca^2+^ out of the cells, thereby leading to the increase of Ca^2+^ concentration in the cytoplasm of sterile pollen, and ultimately leading to pollen abortion.

According to the above DE-TFs, DEGs and DSGs analysis, we found many DEGs annotated as polygalacturonase, glycosyl transferase family 8, ABC transporter were all significantly down-regulated. In addition, DSGs encoding MYB-related protein, ABC transporter B family member 29, pectinesterase 31 and UTP--glucose-1-phosphate uridylyltransferase were identified, which may participate in plant cell wall synthesis (Schubert et al., 2019). In plants, MYB transcription factors play a very important role in the pollen development process, involving various key steps in the pollen formation process, including the programmed cell death (PCD) of tapetum, the deposition of callose, the formation of pollen wall and the accumulation of sporopollenin. Once one of the above abnormalities occurs, it will lead to pollen abortion (Schubert et al., 2019). Previous studies have shown that the large superfamily ABC transporter proteins are involved in translocation of a broad range of substances across membranes using energy from ATP hydrolysis such as transport of sporopollen (the main component of pollen exine) so they are also required for pollen exine formation (Chang et al., 2018). The tapetum provides sucrose, proteins, lipids, and sporopollenin to support the growth and development of the pollen via its degradation and secretion, and the progress of secreting nutrients to pollen is considered to be vesicular trafficking (Liu et al., 2020). Here, vesicular trafficking related DEGs encoding Sec23/Sec24 trunk domain protein, coatomer WD associated region were significantly down-regulated, and identified 15 related DSGs (Fig. 5A and E). To verify the correctness of the above analysis, TEM was used to observe the degradation of tapetum and the formation of pollen wall of BS366 in fertile and sterile conditions (Fig. 9C). During pollen mother cell stage and dyad stage, there was no obvious difference between tapetum of BS366 under two conditions. Up to the tetrad stage, the fertile tapetum structure was complete and connected closely with the middle layer, whereas the tapetum of sterile condition separated from the middle layer. From the release of microspores from tetrapods to the trinuclear stage, the degradation of sterile tapetum was significantly faster than that of fertile tapetum. During this process, the tapetosome of sterile condition were also deformed and missing. Because of the degradation of tapetum in advance, the callose around the tetrad deposited abnormally, subsequently, the pollen wall of microspore released from the tetrad also deformed, which was manifested as the abnormal accumulation and uneven distribution of sporopollenin in the outer wall of pollen. Thus, we conclude that low temperature stress may induce the change of genes encoding MYB transcription factor, ABC transporter and glucose metabolism related enzymes, which may lead to the advanced degradation of tapetum and the deformity of pollen wall.

## Discussion

### Disorder of cytoskeleton is an essential factor for the pollen abortion regulation

In the past decades, many studies showed that cytoskeleton was involved in male sterility in plants. In the PTGMS rice line Peiai 64S displayed abnormal distribution in microtubules at the meiosis stage, no polar microtubules in pollen cells at the zygotene stage and rarefied perinuclear microtubules in diakinesis (Xu et al., 2001). Although Wang et al. (2018) found that disordered and asymmetrical distribution of microflaments and microtubules in sterility pollens, nevertheless, the relationship of cytoskeleton homeostasis with pollen abortion are still unclear in PTGMS wheat line BS366 (Wang et al., 2018). Some studies showed that microtubules display dynamic instability, bouts of rapid growth followed by catastrophic shrinking, and the balance between these phases can be modulated by MAPs which generally increase polymerisation (Kawamura and Wasteneys, 2008). In our study, a MAPs-related hub gene (*TraesCS2D02G523000*), encoding the microtubule-associated protein RP/EB family member 3, was significantly up-regulated during the three stages in the sterile condition in BS366 (Fig. 5E), suggesting that the abnormal expression of MAPs-related gene may lead to the aggregation disorder of cytoskeleton. For our DSGs analysis, 17 DSGs including two genes annotated as *MOR1* were found to be related to cytoskeleton (Table S6). In *Arabidopsis thaliana*, *MOR1*, the homologue of Xenopus *MAP215*, promoted rapid growth and shrinkage, and suppressed the pausing of microtubules in vivo (Kawamura and Wasteneys, 2008). In this study, *MOR1* with differential alternative splicing, may hinder the rapid growth of cytoskeleton abnormally in the critical stage of abortion, thereby affect the development of pollen. In addition, lncRNAs with their targets were also involved in cytoskeleton. Cell cycle regulation related gene *NSA2* (Nop seven-associated 2) could blocked the cell cycle in G1/S transition in *Arabidopsis thaliana*. Here, *NSA2* (*TraesCS2B02G131900*) (target of lncRNA PB.3805.1) was up regulated in SS3 and down regulated in FS3. At the same time, our cytological observation also verified the above analysis (Fig. 9).

### The abnormal calcium messenger system might be related to pollen abortion

At present, it is known that the calcium messenger system is often in the center of signal cascade. Tian et al. (1998) described the anomalies in the distribution of calcium in anthers of PGMS rice, which displayed the failure of pollen development and pollen abortion. It was found that calcium precipitates were abundant in the middle layer and endothecium in sterile anthers, but not in the tapetum (Tian et al., 1998). In this study, calcium messenger system-related genes were excavated from DSG and DEG sets, for instance, *TraesCS6D02G176800*, which was annotated as *BON 1; TraesCS4A02G407100*, which was annotated as Calmodulin-binding transcription activator 2 (*CAMTA*); *TraesCS6B02G230900*, which was annotated as Calmodulin binding protein-like (*CBPL*); *TraesCS5B02G160500*, which was annotated as calmodulin binding receptor like cytoplasmic kinase 3 (*CRCK3*), and *TraesCS7A02G553100*, which was annotated as cytoskeletal-regulatory complex EF hand from DEGs, suggesting that calcium messenger system related genes might play essential roles in pollen fertility transformation after encountering low temperature during meiosis (Table S6). Ca^2 +^, Calmodulin (CaM) and CaM binding propeins (CaMBP) are involved in the process of dynamic distribution of cytoskeleton that regulated b microtubule/microfilement associated proteins (MAPs). Microfilament assembly will be promoted when the concentration of [Ca^2+^]_cyt_ is decreased, and that will be inhibited when the concentration of [Ca^2+^]_cyt_ is increased (Helper and Callaham, 1987). In resistant cowpea, elevation of [Ca^2+^]_cyt_ lead to deassembly of microtubule during rust fungal infection (Xu and Heath, 1998). To future investigated the mechanism of calcium in pollen fertility transformation, the potassium antimonate was used to locate Ca^2+^ in fertile and sterile anthers of BS366. It was found that pollen cell of BS366 with high concentration of [Ca^2+^]_cyt_ and low Ca^2+^ on the cell surface showed pollen abortion, suggested that the reason for pollen abortion may be due to the calcium pump or calcium channel dysfunction, which could not discharge excess Ca^2+^ outside the cell, so that the middle layer of anther was abundant in calcium precipitation (Fig. 9). The up-regulation of calcium-related DEGs and the regulation of DAS may be the reason of calcium message system dysfunction (Fig. 5A and E).

### The abnormal vesicle trafficking is related to pollen abortion

In plants, vesicle trafficking is the main way of material and information exchange between organelles and is also important for maintaining homeostasis, which is involved in many biological processes such as cell wall formation, cell secretion and environmental response (Singh et al., 2018). The membrane vesicle transport machinery includes phospholipids and integral membrane proteins, such as vesicle-associated membrane proteins (VAMPs), the latter being the major constituent of soluble NSF (N-ethylmaleimide-sensitive factor) attachment protein receptors (SNARE) complexes (Han et al., 2017). The vesicle-associated proteins (VAPs) are type II integral ER membrane-bound proteins tethered to the membranes and have been implicated in different processes such as membrane trafficking, lipid transport and metabolism, and unfolded protein response. SNARE complexes are responsible for fusion of vesicles with the target membranes, which is involved in many processes, such as cell plate formation, ion channel regulation, plant growth and development, plant tropism response (Jena, 2011). Sec23/Sec24, the core component of the coat protein complex II (COPII), functions to transport newly synthesized proteins and lipids from the endoplasmic reticulum (ER) to the Golgi apparatus in cells for secretion (Jing et al., 2019). However, information on the role of vesicle trafficking related proteins in wheat pollen development is scanty. In the present study, the gene encoding putative vesicle-associated protein 4-2 (VAP4-2) exhibited significant AS pattern change in PTGMS line BS366. Moreover, the WGCNA analysis results showed the hub DEGs include a soluble NSF attachment protein (SNAP), a conserved oligomeric Golgi complex component (COG2), a dynamin-related protein 5A and three Sec23/Sec24 trunk domain proteins, and it may affect the process of pollen obtaining nutrients through abnormal vesicle transport. Thus, combined with cytological observation and sequencing analysis, we thought that the changes of these genes might result in the abnormal development of pollen. It was also found some DSGs encoding MYB-related protein, UTP--glucose-1-phosphate uridylyltransferase, and ABC transporter B family member 29, as well as some DEGs encoding cytochrome P450 and ABC transporter F family member 3, and these abnormal changes may affect sporopollenin synthesis and formation of the pollen wall (Chang et al., 2016).

### Putative cytoskeleton-related and AS mediated pollen sterile network in PTGMS wheat

According to the putative functions and changes in the DSGs, DEGs, DE-TFs, and lncRNA and their experimental verification in the present study, we propose an intriguing cytoskeleton related transcriptome and AS response mediated pollen sterile network for PTGMS wheat, as shown in Fig. 10. This network has several functional components comprising the calcium regulation, vesicle trafficking, distribution of cytoskeleton and pollen development. The abnormal alternative splicing of these genes encoding kinesin-related protein and myosin as well as the down-regulated genes encoding dynein may hinder the post-translational modification of microtubules, which may further lead to the disordered distribution of microtubules in pollen. Therefore, the low temperature environment, as a signal, may activate or repress the transcription factors, lncRNA or splicing factors of Ca^2+^ and vesicle trafficking, and these in turn regulate the transcription or AS of downstream genes, which in turn disrupted the distribution of the cytoskeleton, thereby hindering pollen development, and ultimately leading to male sterility in BS366.

**Figure 10:**
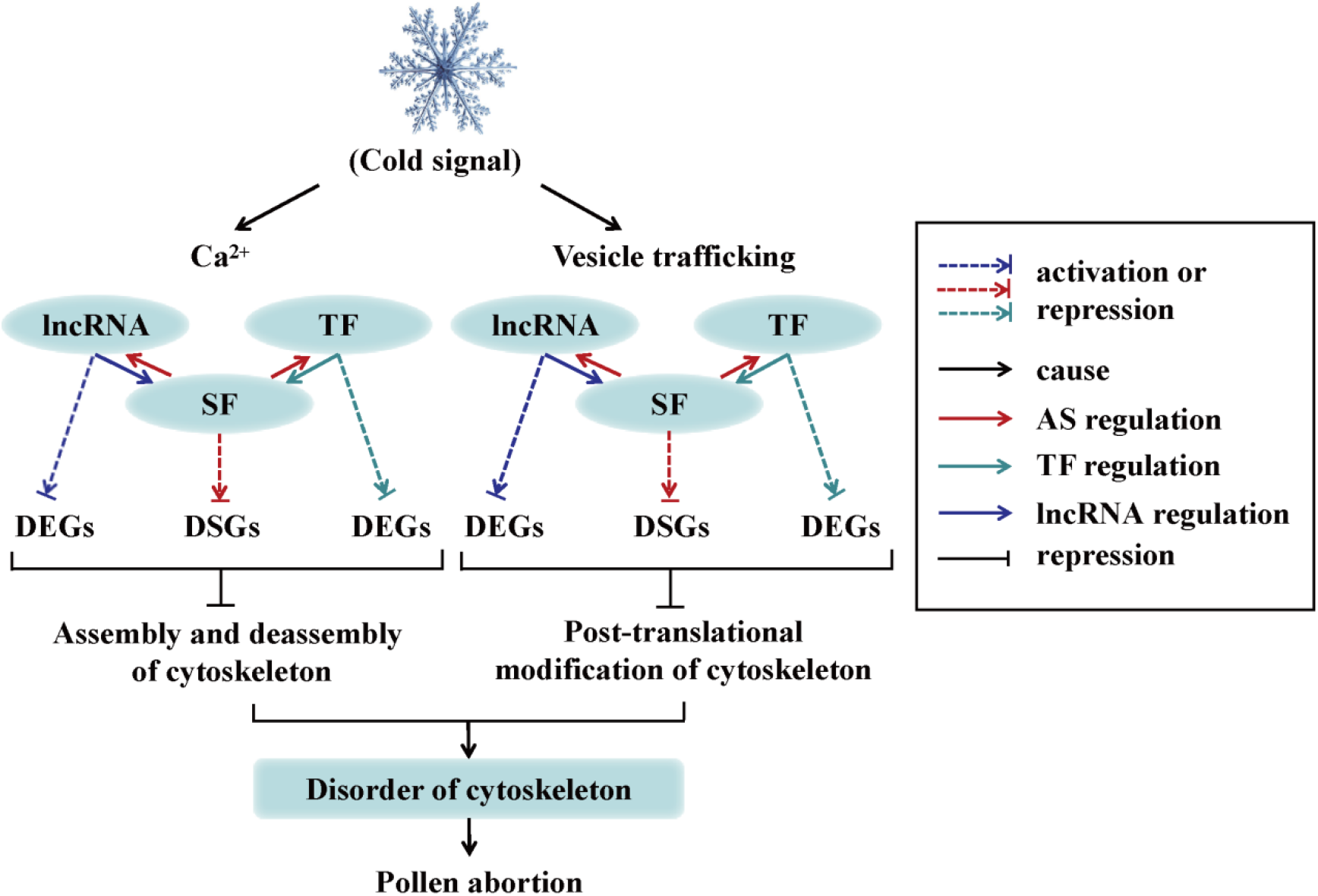
Proposed a cytoskeleton related transcriptome and AS response mediated regulation networks and the signaling pathway involved in male sterility of PTGMS wheat line BS366. Low temperature activates or repress transcription factors, lncRNA or splicing factors of Ca^2+^ and vesicle trafficking and these in turn regulate the transcription or AS of downstream genes, which in turn disrupted the distribution of the cytoskeleton, thereby hindering pollen development, and ultimately leading to male sterility in wheat PTGMS line BS366. lncRNA, long non cording RNA; TF, transcription factor; SF, splicing factors; DEG, differentially expressed gene; DSG, differentially spliced gene.

## Conclusions

In conclusion, through the mechanistic study of the pollen sterility phenotypic change in wheat PTGMS line BS366 by analysis combining second- and third-generation sequencing and investigations of ultrastructural, we demonstrated that Ca^2+^ and vesicle trafficking related DEGs, DSGs and lncRNA affected the assembly and deassembly as well as post-translational modification of cytoskeleton, thereby causing disorder of cytoskeleton, eventually led to pollen sterility (Fig. 10). Our study sheds new light on the underlying mechanism of how cytoskeleton contributes to male sterility in plants and the data could be used as a benchmark for future studies of the molecular mechanisms of PTGMS in other crops.

## Materials and methods

### Plant materials, growth conditions, and sample collection

In this study, the wheat (*Triticum aestivum* L.) PTGMS line BS366 (Bai et al. 2017) and the conventional wheat line Jing411 were used as plant materials. All plants were planted in experimental fields in Beijing (China, N 39°54′, E 116°18′) in plastic pots in early October and managed conventionally. The treatments for wheats and anther sample collection were performed according to Bai et al. (2017). The overall anther development period was divided into six stages: S1: pollen mother cell stage; S2: dyad stage; S3: tetrad stage; S4: uninucleate stage; S5: binucleate stage and S6: trinucleate stage as defined in Browne et al. 2018 (Browne et al., 2018). Samples from fertile and sterile conditions, were collected and named FS1, FS2, FS3, FS4, FS5, FS6 and SS1, SS2, SS3, SS4, SS5, SS6 for each developmental stage, respectively. Of which S1, S2 and S3 were for sequencing, S4, S5 and S6 were for phenotypic characterization assistant.

### Phenotypic characterization at the trinucleate stage

Anthers at the trinucleate stage from fertile and sterile conditions were photographed with a Nikon E995 digital camera (Nikon, Japan) mounted on a Motic K400 dissecting microscope (Preiser Scientific, Louisville, KY, USA). To further analyze pollen fertility, the mature pollen grains were stained using I2-KI staining and photographed with a microscope (Zeiss stemi 305). For SEM analysis, anthers in the trinucleate stage were collected, fixed in 2.5% glutaraldehyde, dehydrated, air dried in silica, coated with gold-platinum in a sputter coater, and finally examined by SEM (Hitachi S-3400N) (Yang et al., 2018).

### RNA Sequencing

After confirming the anther development periods, three critical pollen fertility transformation stages (S1, S2 and S3) were selected for further sequencing analysis (Figure 1). Illumina RNA seq and Iso-Seq library were constructed using the method of Wang et al. (2019). The RNAs of 36 samples (three stages of two cultivars in two different conditions, three biological replicates per stage of two cultivars) were subjected to 150bp paried-end sequencing using HiSeq X Ten platform (Illumina), and then the RNAs of 18 samples from each cultivars were mixed in equal concentration and sequenced on the PacBio RS II platform. Sequencing were performed according to the manufacturer’s standard protocol.

### Identification of full length transcripts

Raw data obtained from Illumina sequencing were processed and filtered by Illumina pipeline (https://www.illumina.com/) to generated FastQ files. Raw data obtained from PacBio sequencing were processed using SMRT Pipe analysis workflow of the PacBio SMRT Analysis software suite (https://www.pacb.com/products-and-services/analytical-software/smrt-analysis/). Raw polymerase reads were filtered and trimmed to generate the ik, subreads and read of inserts (ROIs), requiring a minimum polymerase read length of 50 bp, a minimal read score of 0.65, a minimum subread length of 50 bp, Iso-seq pipeline with minFullPass of 0 and a minimum predicted accuracy of 0.8. Next, full-length, non-chemiric (FLNC) transcripts were determined by searching for the polyA tail signal and the 5’ and 3’ cDNA primers in ROIs. ICE (Iterative Clustering for Error Correction) was used to obtain consensus isoforms and full-length (FL) consensus sequences from ICE was polished using Quiver. High quality FL transcripts were classified with the criteria post-correction accuracy above 99%. Then, FL consensus sequences were mapped to reference genome using Genomic Mapping and Alignment Program (GMAP) using parameters ‘cross-species-allow-close-indels0’ and filtered for 99% alignment coverage and 85% alignment identity (Wu and Watanabe, 2005). Here, 5’ difference was not considered when collapsing redundant transcripts. Integrity assessment for transcripts with no redundant using BUSCO (Simao et al., 2015).

### Fusion transcript delectation

The fusion candidates were detected using criteria that: a single transcript must: 1) map to 2 or more loci, 2) minimum coverage for each loci is 5% and minimum coverage in bp is at least 1 bp, and total coverage is at least 95% and 3) distance between the loci is at least 10kb.

### AS detection and fertility related AS event identification

Transcripts were validated against known reference transcript annotations with the python library MatchAnnot. In this study, AS events including ES, IR, A5’SS, A3’SS and MEX were detected and quantified using rMATS and AStalavista tool (version 3.0) (http://astalavista.sammeth.net/) (Foissac and Sammeth, 2007). Candidate splicing event was calculated using reads mapped to splicing junctions. Differential splicing genes under FS1, FS2, and FS3 compared with SS1, SS2, and SS3, respectively, were selected with FDR ≤ 0.05.

### Identification of lncRNA

The full-length transcripts were aligned to genome of *Triticum aestivum* L. from *Ensembl Plants* database (http://plant.ensembl.org/index.html). Those which could not be aligned were considered as novel transcripts. The novel transcripts (>200bp) were processed to identify lncRNAs based on four computational approaches include Coding Potential Calculator (CPC) (Kong et al., 2007), Coding-Non-Coding Index (CNCI) (Sun et al., 2013), Coding Potential Assessment Tool (CPAT) (Wang et al., 2013) and Pfam. The lncRNA from intersection of these four computational approaches were used further analysis.

### Identification of the transcription factors

The TFs were identified based on the domains of known TFs in the plant transcription factor database PlnTFDB 3.0 (http://plntfdb.bio.uni-potsdam.de/v3.0/). The domains of the protein corresponding to the newly identified transcripts in our analysis and the annotated transcripts in the IWGSC RefSeq v1.0 (generated from both high-confidence genes and low confidence genes) were searched against the included domains and excluded domains of each TF in the PlnTFDB database using the hmm search function of the HMMER software, and only proteins with exactly the same included domains and not with the excluded domains were regarded as TFs. All TFs were against DEGs to confirm the fertility-related TFs.

### Cytological observation

For the transmission electron microscopy (TEM) observation, anthers were fixed, embedded, and stained as described by Zhang et al. (2014). The ultrathin sections were observed and obtained with transmission electron microscope (Hitachi, H-7650, Tokyo, Japan) and an 832 charge-coupled device camera (Gatan, Abingdon, VA, USA). In pollen cells, microfilaments and microtubules were marked by tetramethylrhodamine isothiocyanate (TRITC)-phalloidin (Sigma, St. Louis, MO, USA) and anti-α-tubulin (mouse IgG monoclonal anti-α-tubulin, T-9026; Sigma), respectively. The staining procedures were the same as those described by Wang et al. (2018). For DNA staining, 4’, 6-diamidino-2-phenylindole (DAPI) was used for counterstaining. The DAPI staining procedures were the same as those described by Li et al. (2019). Preparations were observed and images were captured using a laser scanning confocal microscope (Nikon A1R, Tokyo, Japan).

## Data access

The data reported in this article have been deposited in the National Genomics Data Center (NGDC) Genome Sequence Archive (GSA) database under the BioProject accession no PRJCA002516. (https://bigd.big.ac.cn/).Dr

## Acknowledgments

We are grateful to Dr Feng Xu for his helpful suggestions and bioinformatics analysis assistance.

## Figure and table Legends

**Figure S1:** Flowchart of sample collection and RNA-sequencing analysis. The anthers with three stages including pollen mother cell stage, dyad and tetrad stage, were taken from middle of spikes from two condition in both BS366 and Jing411. In total, 36 samples (three stages for each of the two varieties in two conditions, three biological replicates per stage) were sequenced using second-generation sequencing, and two mixed samples (the RNAs of 18 samples from each variety mixed in equal volume) were sequenced using third-generation sequencing

**Figure S2:** Summary of the direct RNA sequencing data of BS366 and Jing411. The bubble scatter plots show the relationship between the fraction of detected transcripts by the direct RNA sequencing with the transcript length and the level transcript expression. The violin-boxplots on the right show the overall distribution of the expression of transcripts.

**Table S1:** Primers used in this study

**Table S2:** Summary information of circular consensus sequence reads

**Table S3:** Summary Statistics of Full-length non-chimeric and isform

**Table S4:** Summary Statistics of transcript and gene loci

**Table S5:** Statistics of alternative splicing events in different anther development stages during fertility transition in BS366 and Jing411

**Table S6:** GO analysis of pollen sterile related DAS, DSGs and targets of lncRNA

**Table S7:** The seed setting rate of BS366 and Jing 411 in different conditions

**Table S8:** Identificatied TFs from Pacbio data

**Table S9:** Statistics of differentially expressed TFs (DE-TFs) and differentially spliced TFs (DS-TFs) in different anther development stages

**Table S10:** lncRNA and their corresponding targets

**Table S11:** Annotation of targets of lncRNA

